# Signal-to-Signal Neural Networks for Improved Spike Estimation from Calcium Imaging Data

**DOI:** 10.1101/2020.05.01.071993

**Authors:** Jilt Sebastian, Mriganka Sur, Hema A. Murthy, Mathew Magimai.-Doss

## Abstract

Spiking information of individual neurons is essential for functional and behavioral analysis in neuroscience research. Calcium imaging techniques are generally employed to obtain activities of neuronal populations. However, these techniques result in slowly-varying fluorescence signals with low temporal resolution. Estimating the temporal positions of the neuronal action potentials from these signals is a challenging problem. In the literature, several generative model-based and data-driven algorithms have been studied with varied levels of success. This article proposes a neural network-based signal-to-signal conversion approach, where it takes as input raw-fluorescence signal and learns to estimate the spike information in an end-to-end fashion. Theoretically, the proposed approach formulates the spike estimation as a single channel source separation problem with unknown mixing conditions. The source corresponding to the action potentials at a lower resolution is estimated at the output. Experimental studies on the spikefinder challenge dataset show that the proposed signal-to-signal conversion approach significantly outperforms state-of-the-art-methods in terms of Pearson’s correlation coefficient and Spearman’s rank correlation coefficient and yields comparable performance for the area under the receiver operating characteristics measure. We also show that the resulting system: (a) has low complexity with respect to existing supervised approaches and is reproducible; (b) is layer-wise interpretable; and (c) has the capability to generalize across different calcium indicators.

**Author summary:** Information processing by a population of neurons is studied using two-photon calcium imaging techniques. A neuronal spike results in an increased intracellular calcium concentration. Fluorescent calcium indicators change their brightness upon a change in the calcium concentration, and this change is captured in the imaging technique. The task of estimating the actual spike positions from the brightness variations is formally referred to as spike estimation. Several signal processing and machine learning-based algorithms have been proposed in the past to solve this problem. However, the task is still far from being solved. Here we present a novel neural network-based data-driven algorithm for spike estimation. Our method takes the fluorescence recording as the input and synthesizes the spike information signal, which is well-correlated with the actual spike positions. Our method outperforms state-of-the-art methods on standard evaluation framework. We further analyze different components of the model and discuss its benefits.

## 1 Introduction

Behavioral and other cognitive aspects of the brain are analyzed based on the brain’s responses to several types of stimuli. The brain responds in the form of spikes, which digitally encodes the information. The analyzes can be carried out at the neuronal-level (micro) or over a brain area covering several thousands of neurons (macro-level). The behavior of individual neurons across a specified time is analyzed in micro-level in comparison to the macro-level analyzes, where the pattern of activation across a population of neurons is considered. The latest scanning methods track the activity of a population of neurons by using fluorescence emitting capability of calcium indicator proteins/dye [1–4]. However, the calcium fluorescence recording of each neuron is only an indirect indicator of the actual spiking process. The presence of fluorescence level fluctuations, slow dynamics of the calcium fluorescence signal, and unknown noise-levels make it hard to identify the exact underlying spike information [5–7]. Hence, technologies capable of obtaining the spike positions from the calcium fluorescence signals are of utmost interest to the computational neuroscience community.

Existing spike estimation algorithms can be broadly categorized into generative and data-driven approaches:

1. Generative methods model the fluorescence signal as the response of the calcium indicator to the spike occurrences. They rely on several model-specific assumptions. Deconvolution-based approaches consider convolutive assumptions about the spiking process [8–10, 38], whereas biophysical model-based approaches estimate the most probable spike train which generated the fluorescence output [11]. Other model-based approaches include template matching [3, 12], auto-regressive formulation [32] and, approximate Bayesian inference based on deconvolution [5, 13]. These models are limited by the apriori assumptions about the model, which has stringent approximations regarding the shape of the calcium response and the noise statistics. Recently, a non-model based signal processing approach for spike estimation is presented in [16] which uses the agnostic nature of group delay to estimate the spike locations. It has a comparable performance with other popular algorithms in the literature.
2. Supervised models predict the spike information from the fluorescence signal, either using a set of features derived from the signal or using the raw signal itself. Data-driven methods are recently gaining traction owing to the availability of simultaneous electrophysiological and two-photon scanning-based neuronal recordings. For instance, a neural network-based supervised Spike Triggered Mixture (STM) model [17] is used for learning the *λ* parameter of a given Poisson model in [18] to obtain the spike estimates from the calcium signals. Recent methods use fluorescence signals with or without a contextual window (supplementary material - [15]) for estimating the spike information. Neural network-based variants such as “convi6”, “Deepspike”, “Purgatorio’, and “Embedding of CNNs” have had varying levels of success and outperformed data-driven baseline method [17] on a standard evaluation framework (supplementary material - [15]). A gated recurrent unit (GRU)-based approach recently attempted to estimate down-sampled action potentials directly from the 2-D calcium imaging output, combining regions-of-interest (ROI) estimation and spike estimation tasks [19]. An adversarial variational autoencoder (VAE) is employed in [14] for improved spike inference compared to the factorized posteriors used in standard VAEs.

Spikefinder challenge (http://spikefinder.codeneuro.org/) [15] resulted in a new set of algorithms that performed better than the benchmark STM Model [17]. Although these algorithms use different techniques, it was found that their fusion could not improve the results further. The challenge was aimed to standardize the spike estimation evaluation and to provide a comparison of the state-of-the-art techniques. Most of the top-performing algorithms used convolutional (CNN), recurrent (RNN) and, deep neural networks (DNN) and its variants [15]. All the top-performing data-driven algorithms have a recurrent layer in the network and hence are computationally complex. The best-performing supervised model used a CNN architecture with an intermediate Long Short-Term Memory (LSTM) layer to predict the spiking probability from a contextual window of the fluorescence signal (“convi6” in the supplementary material of [15]). Generative models such as MLspike [11] and auto-regressive model presented in [32] were comparable to the supervised approaches after dataset-specific parameter tuning carried out on the training datasets. MLspike uses a biophysical model and estimates the maximum probable spike information given the fluorescence signal. The second-best generative approach in spikefinder [32] is based on an autoregressive approximation to the calcium fluorescence signal. Spike information is then estimated by solving a non-negative sparse optimization problem.

The spike estimation problem can be formulated as a sequence-to-sequence modeling problem in the supervised learning paradigm. Here we briefly describe the motivation and related works on sequence modeling to set the stage for the proposed method. Approaches that model sequences in an end-to-end fashion are being developed recently for classification and regression tasks in audio processing and natural language understanding with commendable success. One of the first efforts in that direction was made in the context of machine translation [22]. This approach was later extended to speech recognition [23, 24]. Sequence-to-sequence models have been used for text summarization task using RNNs in [20] and for creating more accurate language models [21] in natural language processing. End-to-end methods have also been employed to predict the target speech from the overlapped speech mixture for speech separation task [25, 26], and clean speech for enhancement tasks [27, 28] in audio processing research. In this case, as the models are trained on raw-waveforms, they do not necessitate handcrafted features. Instead, they learn the relevant information in a task-dependent manner from the signal directly. They also provide the flexibility to choose task-specific objective functions for training, which implicitly considers the temporal context. It is also possible to analyze the convolutional filters in the neural network to understand the learning trends in the temporal and frequency domains. In the same way, it should be possible to formulate the spike estimation problem and analyze the filters to get more insights from them.

Motivated by the success of aforementioned sequence-to-sequence models, we present a signal-to-signal neural network (hereafter referred to as S2S) for the spike estimation task. The proposed method can be regarded as an analysis-synthesis method, where, as illustrated in Fig 1, the input calcium fluorescence signal is analyzed by an input convolution layer; filtered by time-distributed dense layer(s) (also called as hidden layers); ^1^ and finally the spike signal is synthesized by an output convolution layer. The synthesis layer that generates signal samples makes our architecture distinct from other supervised spike estimation methods where only a single value is predicted/classified to at the output. All the network parameters are learned in an end-to-end manner in S2S with a cost function based on the Pearson’s correlation between the estimated spike signal and the ground truth spike signal. We hypothesize that such a spike estimation network can outperform existing approaches as it *reconstructs* the spike information for each input sample. This neural network differs from the sequence-to-sequence models mentioned earlier (Lines 76-81) in several aspects. First, the output signal’s nature or characteristics (discrete spike estimates) is very different from the input signal (calcium fluorescence trace). Second, it performs automatic short-time processing through shared weights across the temporal axis. Third, as we will see later (Section 2.4.4), each layer’s output can be visualized to gain insight into the information captured by each layer. The frequency responses of both analysis and synthesis layers in the network can also be analyzed.

**Fig 1.**
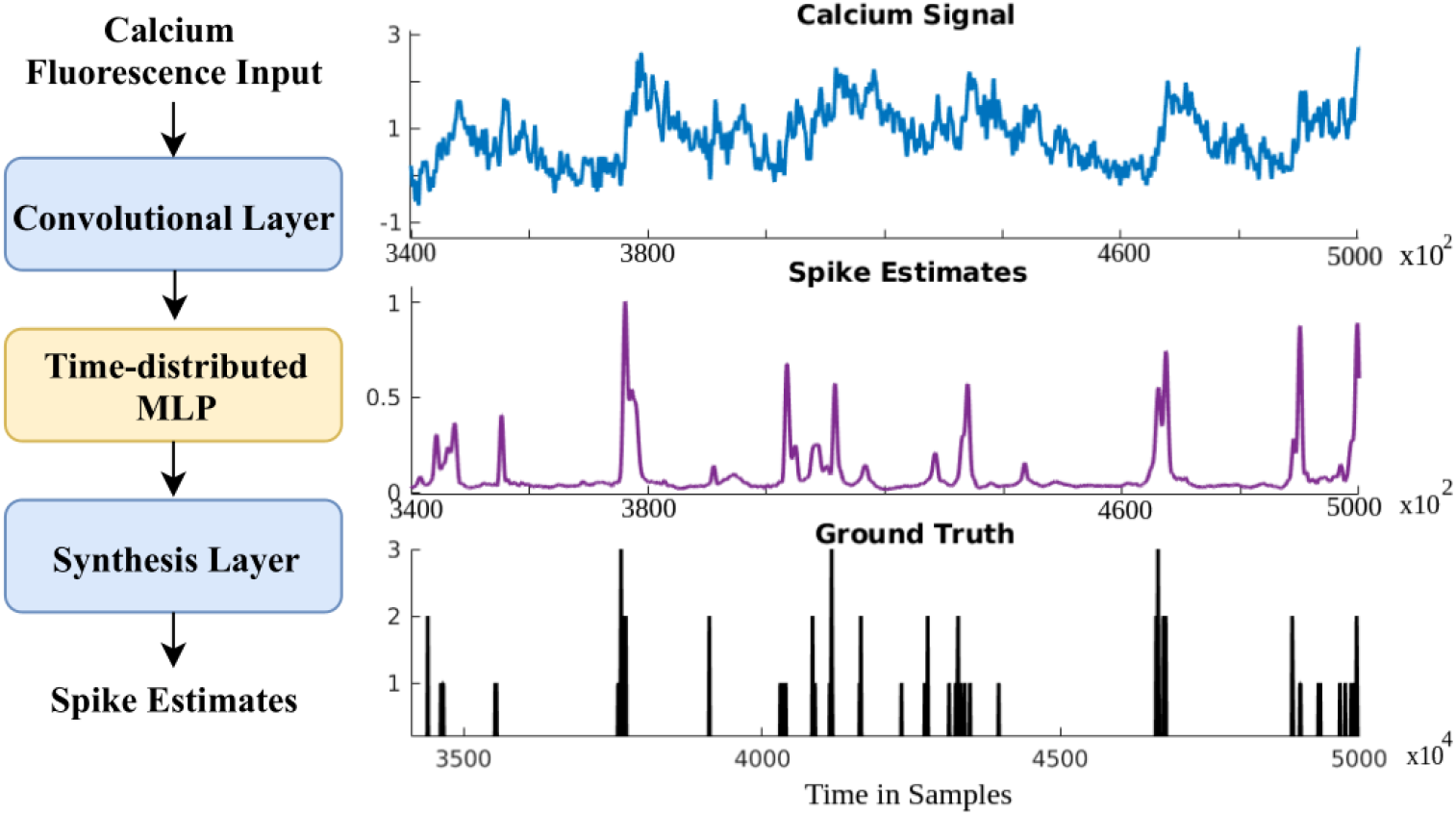
Signal-to-Signal neural network (S2S). (***left***) Block diagram of the proposed approach. (***right***) An illustrative example calcium signal, its corresponding spike estimate, and the ground truth spike train.

The proposed S2S method is compared with competitive algorithms from the spikefinder contest with the same dataset and evaluation procedure. A detailed comparison of S2S is also made with the best neural network-based model presented in the contest. The research questions in terms of reliability, generalization ability, dependency on training targets and, design concerns of S2S are studied. We also conducted a layer-wise analysis of the network to provide an intuitive explanation for the learning.

## 2 Results

### 2.1 Evaluation Procedure

We used the Spikefinder challenge dataset [15] for evaluations. It consisted of five benchmarking datasets consisting of 92 recordings from 73 neurons. One part of this dataset was given for training the supervised models and the other part for testing as a part of the competition. Five datasets [30] from GENIE project [41] were also available for training the models. These additional datasets were to make sure that the supervised models do not over-fit to the training data. For further details about the spike finder dataset, the reader is referred to [15]. As per the spike finder challenge protocol, we performed the training at 100 Hz and the evaluation at 25 Hz (40 ms bin width). The evaluation measures were Pearson’s correlation coefficient, Spearman’s rank correlation coefficient, and the area under the receiver operating characteristics (denoted as AUC).

We used the Pearson’s correlation coefficient as the primary evaluation measure, as done in the spikefinder challenge. Rank (non-linear correlation) and AUC serves as secondary and tertiary evaluation measures, respectively. This standardization enabled us to benchmark the proposed S2S method against the challenge submissions. It is worth mentioning that the performance metrics were calculated solely based on the script provided by the spikefinder challenge organizers. The performance of the S2S method is compared to the top six algorithms in the spikefinder contest. They were either based on generative [11, 32] or supervised [15, 17, 33, 39] approaches, as discussed earlier (Lines 18-48). Seven out of top-10 algorithms were deep learning-based supervised algorithms. Table 1 provides an overview of the baseline methods (taken from [15]).

**Table 1.** Overview of top-performing algorithms in spikefinder challenge.

| Team | Contributor(s) | new | type | Model/Architecture |
| --- | --- | --- | --- | --- |
| Team1 | T. Deneux | No [11] | Generative | Biophysical model |
| Team2 | N. Chenkov, T. McColgan | Yes | Supervised | conv / lstm |
| Team3 | A. Speiser, J. Macke, S. Turaga | Yes | Supervised | RNN/CNN |
| Team4 | P. Mineault | Yes | Supervised | residual / lstm |
| Team5 | P. Rupprecht, S. Gerhard, R. W. Friedrich | Yes | Supervised | conv / max |
| Team6 | J. Friedrich, L. Paninski | No [32] | Generative | Autoregressive model |
| Baseline: STM | L. Theis | No [17] | Supervised | DNN +Poisson model |

### 2.2 Comparison to Spikefinder Algorithms

Table 2 compares the performance of the S2S method with the methods reported in the spikefinder challenge [15]. The S2S network outperformed all the state-of-the-art methods, both generative and supervised. It improved the test correlation by 46% compared to the best performing algorithm in the spikefinder contest. Change in the correlation with respect to the challenge baseline (denoted as Δ) is significantly high for the proposed approach (3.5 times compared to the best reference algorithm). S2S provided an improvement of 56% for the rank measure over the baseline with the best rank. It also had a similar AUC compared to the best method in terms of AUC measure. The deviation between the train and test sets’ correlation coefficient for the S2S method was minimum (0.0079). Interestingly, S2S is the only method for which the test set’s correlation coefficient was more than the training set, indicating that the proposed method generalizes well. Furthermore, as presented in Table 3 by numerical values, the performance of the proposed approach over the five different test datasets showed a commendable correlation to the shape and monotonicity of the predicted spike information and ground truth.

**Table 2.**
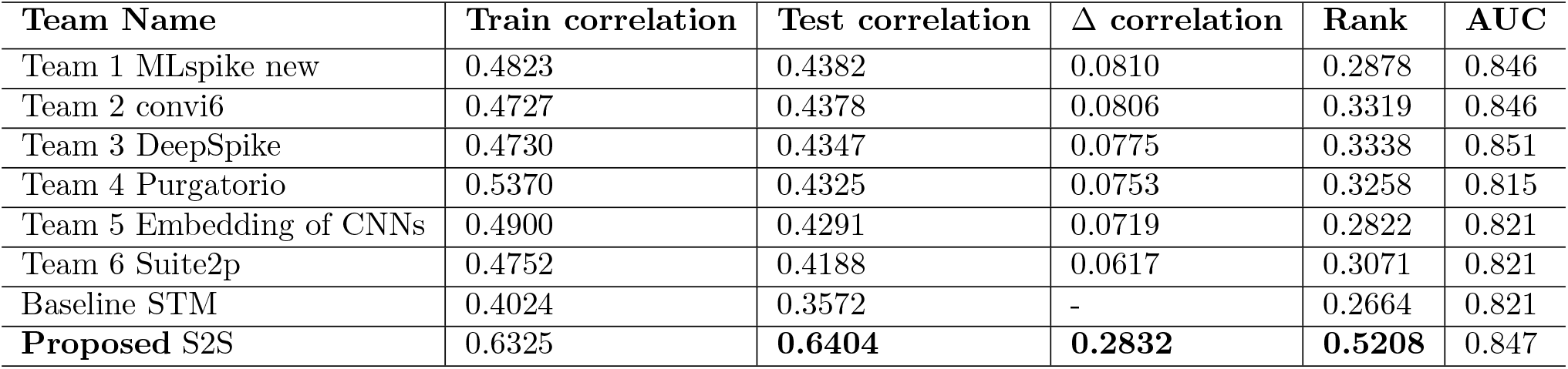
Comparison of evaluation measures between S2S and state-of-the-art spikefinder baselines.

**Table 3.** Dataset-wise performance of S2S on the spikefinder test set.

| Measure | Dataset 1 | Dataset 2 | Dataset 3 | Dataset 4 | Dataset 5 | Average |
| --- | --- | --- | --- | --- | --- | --- |
| Correlation | 0.657 | 0.910 | 0.585 | 0.440 | 0.611 | 0.640 |
| Rank | 0.420 | 0.816 | 0.525 | 0.261 | 0.581 | 0.521 |
| AUC | 0.901 | 0.901 | 0.874 | 0.682 | 0.879 | 0.847 |

Spike trains are recorded originally at a very high sampling rate (10,000 Hz). The Spike train is down-sampled, and the calcium fluorescence signal is up-sampled to 100Hz for the experiments. Spike trains at 25 Hz (40 ms time-duration) can be effectively inferred via S2S, as seen from this evaluation. Since the resolution of the down-sampled spikes is much less than the original frequency, it can be considered as firing rate. S2S is similar to other methods when considering the relative counts of spikes but has a better overall shape of the spike information signal. Better correlation values lead to better sensitivity towards small calcium events that results in a spike (Please refer to the Discussion Section 3.1).

### 2.3 Comparison to State-of-the-Art Supervised Baseline

Fig 2 compares the performance of S2S with the best supervised baseline (Base) “convi6” (Team 2 in Table 2). The baseline approach (convi6) had the second position in the challenge, falling behind MLspike gracefully by 0.04%. S2S achieved significantly better Pearson’s and Spearman’s correlation values across all the five different test sets and achieved comparable AUC. We performed paired t-test on the spikefinder test data to determine the statistical significance of S2S as compared to the convi6 model. Each test dataset was considered as a separate example file similar to that in the spikefinder challenge for the significance test. The null hypothesis (H) was that there is no difference in the performance metrics (correlation, rank and AUC) between S2S and convi6, at 95% confidence level (*α* = 0.05). Table 4 shows the results of the statistical significance test for all of the evaluation measures. The t-value from the table for a degree of freedom 4 is found to be 2.132. The t-value 1 denotes the paired t-test between convi6 and S2S with Gaussian target (Please refer to Section 2.4.1 for details on the training target), and t-value 2 denotes the t-test between convi6 and S2S with the actual training target.

**Fig 2.**
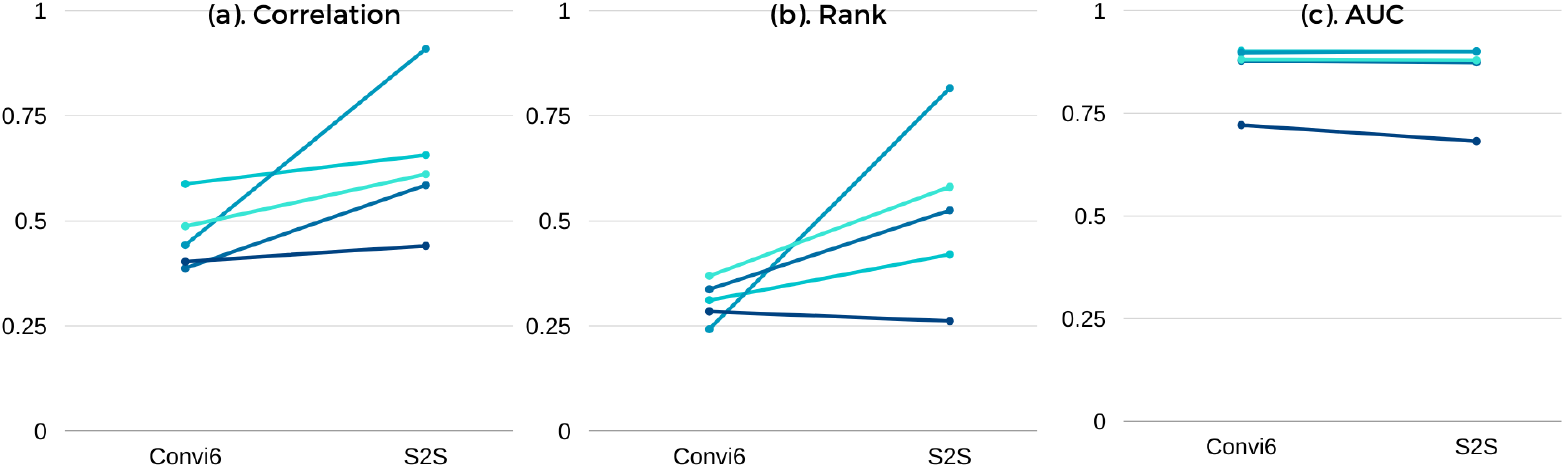
Dataset-wise performance to show the difference in evaluation measures between S2S and convi6. (a) Correlation measure, (b) Rank (non-linear correlation) measure and (c) AUC measure.

**Table 4.** Results on paired t-test for statistical significance. “H” represents the hypothesis that there is no difference in performance between S2S and convi6.

| Measure | t-value 1 | t-value 2 | Accept/Reject H |
| --- | --- | --- | --- |
| Correlation | 2.32 | 2.33 | Reject |
| Rank | 2.14 | 2.16 | Reject |
| AUC | 1.11 | 1.00 | Accept |

There is a significant improvement in the correlation and rank measures and no statistically significant change in the AUC measure. It can also be understood by looking at the mean and standard deviation of the baseline methods. The value of primary evaluation measure (0.640) is greater than the mean and two times the standard deviation (Mean +2× STD) of the baseline approaches (0.479), by 16.1%.

To further validate the scaling ability of S2S at a higher resolution, we evaluated the performance of the same model at 100 Hz (10 ms) sampling rate. The results are shown in Table 5. Correlation measures were affected by the change in the sampling rate. The difference in AUC was negligible from 25 Hz to 100 Hz. We observed that the correlation measures were still better than all other algorithms evaluated at 25 Hz. There was an obvious reduction in performance owing to the higher sampling rate, and it was observed for the baseline algorithms as well. For instance, a considerable amount of reduction in performance was noted for the baseline convi6 model when evaluated at 100 Hz (Please refer to Table 5).

**Table 5.** Results on scalability experiments at 25 Hz and 100 Hz sampling rates.

| Approach | Evaluation Sampling Rate | Correlation | Rank | AUC |
| --- | --- | --- | --- | --- |
| S2S | 25 Hz | 0.640 | 0.522 | 0.848 |
| S2S | 100 Hz | 0.527 | 0.448 | 0.846 |
| Convi6 | 25 Hz | 0.461 | 0.309 | 0.845 |
| Convi6 | 100 Hz | 0.367 | 0.181 | 0.844 |

Signal conversion helped in the reconstruction of spike information that closely resembles discrete ground truth. S2S had reliable output performance corroborated by similar output after a new training process, independent of the parameters’ initialization. We investigated the retraining of the neural networks (new initialization and training on the same dataset) in the S2S method and compared with the convi6 method. We found that the S2S method was insensitive to re-initialization, while convi6 method performance varied on average (standard deviation of 5%, 8%, and 1% absolute for correlation, rank, and AUC, respectively). These results show that the proposed S2S method yield reliable estimates of the spike signal. The best baseline performance (for primary evaluation measure) is observed when using the weights provided by the spikefinder [15]. S2S was significantly faster than the baseline. On Sun GPU clusters, S2S training is 50 times faster than that of the baseline model. Other methods such as MLspike (winner of the spikefinder competition) and Vogelstein are based on signal processing and do not require training. The inference time of the neural network-based methods (convi6, S2S, and others) is *O*(*n*). Both the Vogelstein and STM (test) runs in linear time. MLspike takes *O*(*nlogn*) for dynamic programming and additional time to auto-calibrate its parameters.

### 2.4 Analysis

In this part of the article, we provide analysis of the proposed S2S method in terms of architecture, training target, generalization capabilities and layer-wise interpretation of the trained network.

#### 2.4.1 Training Target

As discussed in the Methods section (Section 4.1.2), the S2S network was trained with a modified ground truth or target by convolving the original discrete spike information with a Gaussian function. Although the Gaussian window hyper-parameters (11, 5), where (*x, y*) denotes Gaussian window width *x* and standard deviation *y* in the number of samples, were obtained through cross-validation during the network training, a question that arises is: what is the impact of less-sparse (more-dense) targets? So, we investigated the S2S network’s training by convolving the target spike signal with the Gaussian windows of variable sizes. For the sake of clarity, Fig 3 compares the performance obtained for (11, 5) with the performance obtained for (33, 11) and the performance with a discrete spike signal as the target signal. It can be observed that convolving with Gaussian function helps. The performance in terms of all three evaluation measures improves from a discrete spike signal to a Gaussian target. Increasing the width of the Gaussian window beyond (33, 11) resulted in reduced performance, as the shape of the training target became very different from the discrete target (actual ground truth). This hyperparameter might be related to the firing rate of the indicators used in the training dataset. We observe that a window of (11, 5) suits best for the spikefinder challenge dataset. There was a little improvement for all of the evaluation measures upon providing a smoothed Gaussian as the target. The difference in performance was 0.1% for the correlation measure in the spikefinder test set and around 12% for the training set. Even with a non-smoothed (“actual”) ground truth, the system’s overall performance is better than the baseline methods (Fig 3 (***left***) and Fig 2).

**Fig 3.**
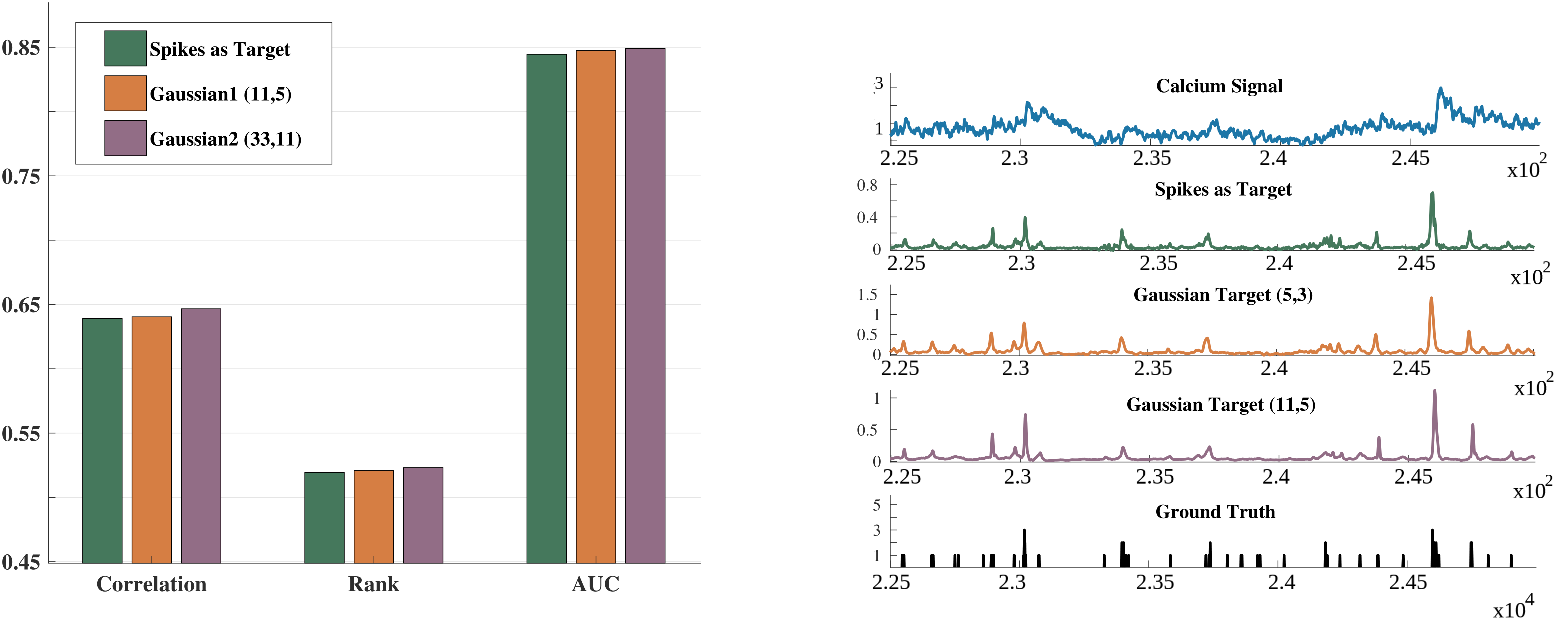
Change in the evaluation measures with Gaussian windowing of the training targets. (***left***) Bar diagram depicting the difference in Correlation, Rank and AUC when using the 100 Hz spike train as the target and when convolving this target with a Gaussian window to generate a smoothed training target. Two different windowing sizes are shown; 11 samples with 5 standard deviation and 33 samples with 11 standard deviation. (***right***) From top to bottom: An example calcium fluorescence signal and its corresponding training targets (spikes at 100 Hz, Gaussian targets with 5 and 11 samples each, and the ground truth at 10,000 Hz). Gaussian training targets are having an equivalent shape to the spikes target at 100 Hz. Note that all of them are approximations of the original ground truth.

In order to further validate the effect of windowing, we performed 10-fold cross-validation of the spikefinder dataset and reported the performance in Table 6. Five training sets and five test sets were used for the leave-one-dataset-out cross-validation. The Gaussian window-based smoothing of the discrete spike target helped improve the correlation and rank measures by 5% and 7.5%, which is a notable difference.

**Table 6.**
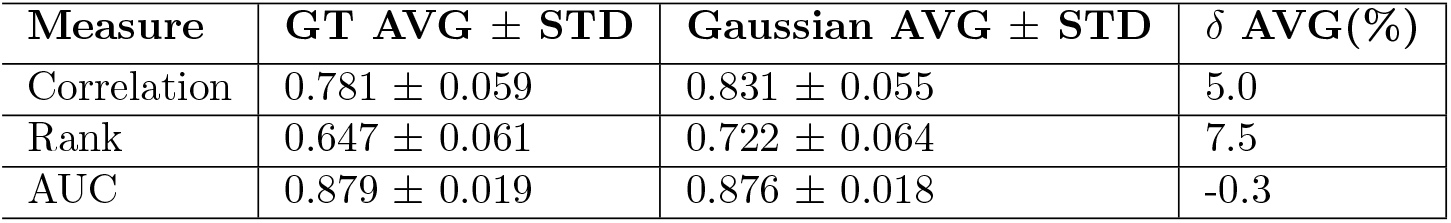
Results on 10-fold cross validation with and without Gaussian windowing for the training target. GT: Ground Truth, AVG: Average and STD: Standard Deviation.

| Measure | GT AVG ± STD | Gaussian AVG ± STD | $\delta$ AVG(%) |
| --- | --- | --- | --- |
| Correlation | 0.781 ± 0.059 | 0.831 ± 0.055 | 5.0 |
| Rank | 0.647 ± 0.061 | 0.722 ± 0.064 | 7.5 |
| AUC | 0.879 ± 0.019 | 0.876 ± 0.018 | -0.3 |

#### 2.4.2 Architecture

The number of hidden layers in the network architecture (explained in the Section 2.4.2) was chosen based on cross-validation. We examined the role of the hidden layers by varying the number of hidden layers from three (used in the experiments reported in the Results part) to zero. The target for training was obtained by convolving the Gaussian window (11, 5) with the discrete spike signal. Fig 4 presents the results in terms of the three evaluation measures. It can be observed that, even without a hidden layer, the proposed S2S method yields a system that is competitive to state-of-the-art systems. The performance improves with the hidden layers. We found that the performance was saturated beyond three hidden layers. As illustrated in the subplot with an example of the input signal, S2S with the hidden layer(s) improves the denoising and spike-resolving capabilities compared to the S2S without a hidden layer.

**Fig 4.**
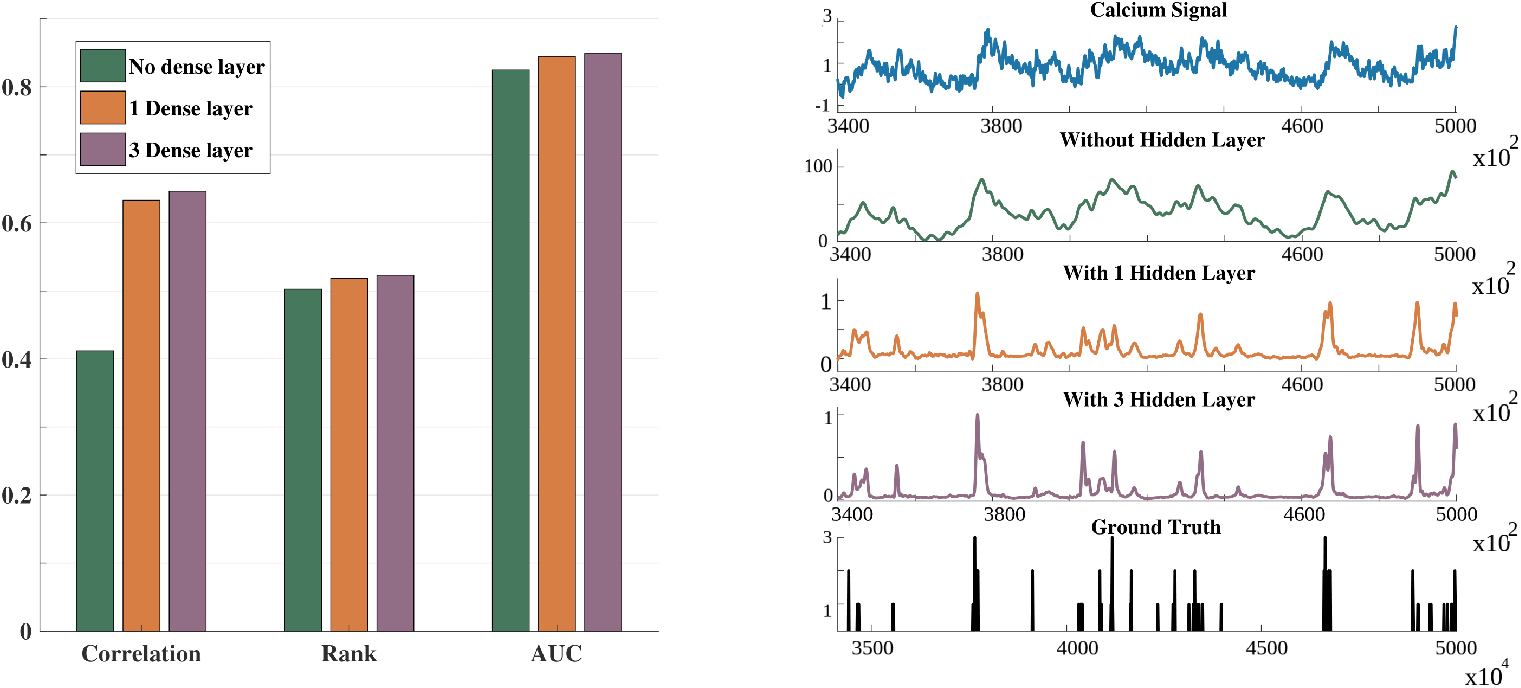
Evaluation measures with changes in number of hidden layers. (***left***) Bar diagram depicting the difference in correlation, rank and AUC when no hidden layer, 1 hidden layer and 3 hidden layers are used in the S2S network, respectively. (***right***) Illustrative example showing the improved spike estimates when 3 hidden layers are used compared to one hidden layer.

#### 2.4.3 Generalization Ability

The spikefinder challenge consists of data obtained with two different indicators, namely, GCaMP indicator and OGB indicator. In the challenge protocol, both the training and test conditions contain signals from these indicators. We examined the proposed approach’s generalization ability across different indicators by training the S2S only on the GCaMP indicator data in the training set and testing on the test set of spikefinder contest containing data from both indicators. More specifically, the *invariance* of the spikefinder test set performance to a change in the training set is considered. We trained S2S with one hidden layer and three hidden layers. The training was carried out with targets obtained by convolving the ground truth discrete spike signals with Gaussian window (11, 5). Table 7 presents the results for both one and three hidden layer S2S. It can be observed that the S2S trained only with the GCaMP indicator yields competitive performance with that trained using all the training data. Furthermore, it can be observed in Fig 5 that the evaluation measures on OGB datasets only gracefully degraded for the GCaMP-only trained model in comparison with the combined-model for *all* the datasets.

**Table 7.** Generalization across indicators. Experimental results with various number of hidden layers.

| Configuration | Training Indicator(s) | Correlation | Rank | AUC |
| --- | --- | --- | --- | --- |
| 1 hidden layer | GCaMP | 0.608 | 0.510 | 0.832 |
| 1 hidden layer | GCaMP + OGB | 0.639 | 0.519 | 0.845 |
| 3 hidden layers | GCaMP | 0.622 | 0.518 | 0.843 |
| 3 hidden layers | GCaMP + OGB | 0.640 | 0.521 | 0.847 |

**Fig 5.**
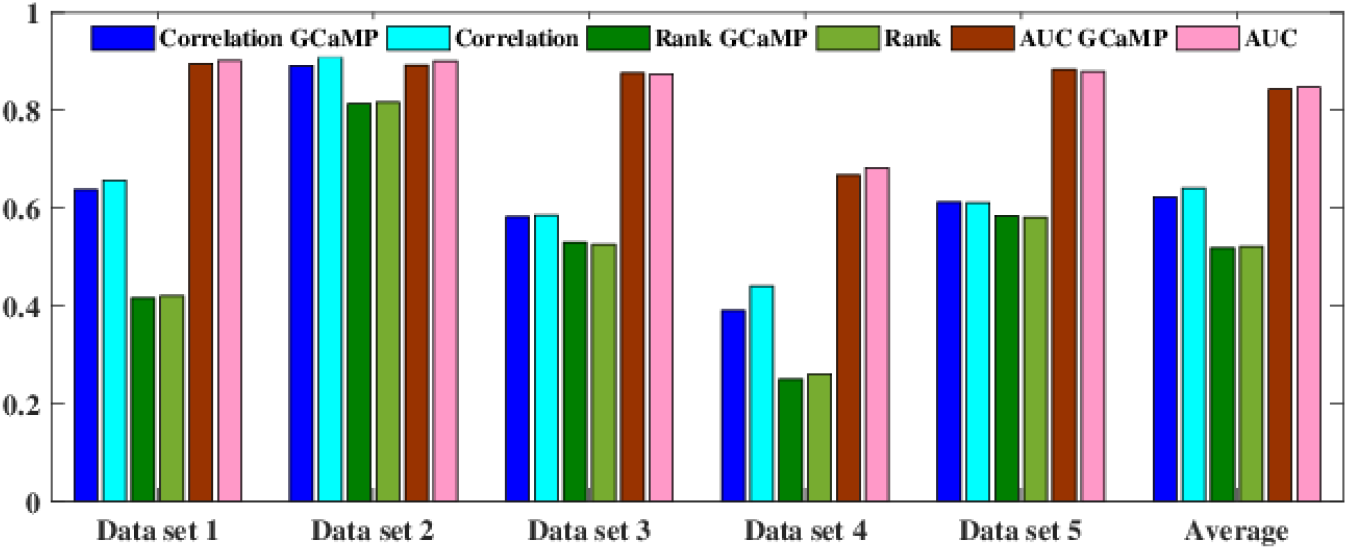
Dataset-wise performance showing the generalization ability of the three hidden layer S2S network. “GCaMP” indicates that the training was done only with GCaMP indicator dataset. Datasets 1, 2 and 4 are based on OGB indicator.

We extended the experiments further by considering only one indicator for the training and the other for testing. For training a GCaMP-only model, we used two training datasets in the spikefinder challenge and the five optional datasets provided for training. For training the OGB-only model, the datasets from both train and test splits of spikefinder challenge were taken. This resulted in 6 OGB datasets for training and 2 GCaMP datasets for testing. The results of the experiments are given in Table 8.

**Table 8.** Results on generalization experiments. Experiments 2b and 3b represents generalization from GCaMP to OGB and vice-versa. For every experiments, the number of datasets used for training and testing are given inside the brackets.

| Sl No. | Experiment | Correlation | Rank | AUC |
| --- | --- | --- | --- | --- |
| 1a | Train full (10) & Test full (5) | 0.640 | 0.522 | 0.848 |
| 1b | Train on GCaMP (7) & Test full (5) | 0.635 | 0.516 | 0.838 |
| 2a | Train full (10) & Test on OGB (3) | 0.672 | 0.498 | 0.827 |
| 2b | Train on GCaMP (7) & Test on OGB (3) | 0.656 | 0.488 | 0.811 |
| 3a | Train full (10) & Test on GCaMP (2) | 0.594 | 0.558 | 0.881 |
| 3b | Train on OGB (6) & Test on GCaMP (2) | 0.465 | 0.493 | 0.803 |

The results corresponding to rows 1a and 1b in Table 8 consider the performance invariance of S2S on the spikefinder challenge test set with a GCaMP-only based training. It can be observed that there is a drop of 0.5% in the primary evaluation measure. These are not significant in comparison to the drop in performance between S2S and the baseline algorithms. Rows 2a and 2b correspond to the differences in OGB performance when only the GCaMP is used for the training. The reduction in performance is 1.6%, 1% and, 1.6%, respectively, for correlation, rank, and AUC. A somewhat different observation can be made for generalization performance for OGB indicators (Rows 3a and 3b in Table 8). The absolute difference in correlation, rank, and AUC are 13%, 6%, and 8%, respectively.

From the results, it can be observed that S2S is generalizable from GCaMP to OGB. However, similar conclusion can not be clearly drawn for the case OGB to GCaMP. A potential reason for that could be the less training data available for OGB-only training.

#### 2.4.4 Layer-wise Output

The S2S method is interpretable, in the sense that the processing carried out by each layer in the network can be visualized to gain insights. Fig 6 shows the layer-wise outputs of the three hidden layer version of the network. These outputs are generated by feeding the calcium input signal and calculating the total response per layer with each layer’s trained weights. For more details regarding the estimation of layer-wise output, the reader is advised to refer to the Methods section (Section 4). For the given calcium fluorescence input, the analysis convolution layer seems to make it less-noisy (or smooth out slight variation in the signal), while preserving possibly the information “relevant” for spike estimation. Observe that the shape of the input signal is preserved. The first hidden layer output after ReLU non-linearity seems to resolve potential spike positions. This output is then refined or filtered by the subsequent hidden layers and the output synthesis layer by enhancing the signal at the spike locations and suppressing the spurious spike locations. This yields an output spike signal estimate that correlates well with the ground truth. Each layer contributes towards maximizing the similarity or correlation between the spike signal estimate and the ground truth. It is worth noting that the output of hidden layers H2 and H3 do not differ too much when compared to H1. The output synthesis convolution layer seems to carry out the significant refinements. This observation is in line with the earlier observation that one hidden layer S2S yields performance comparable to the three hidden layer S2S.

**Fig 6.**
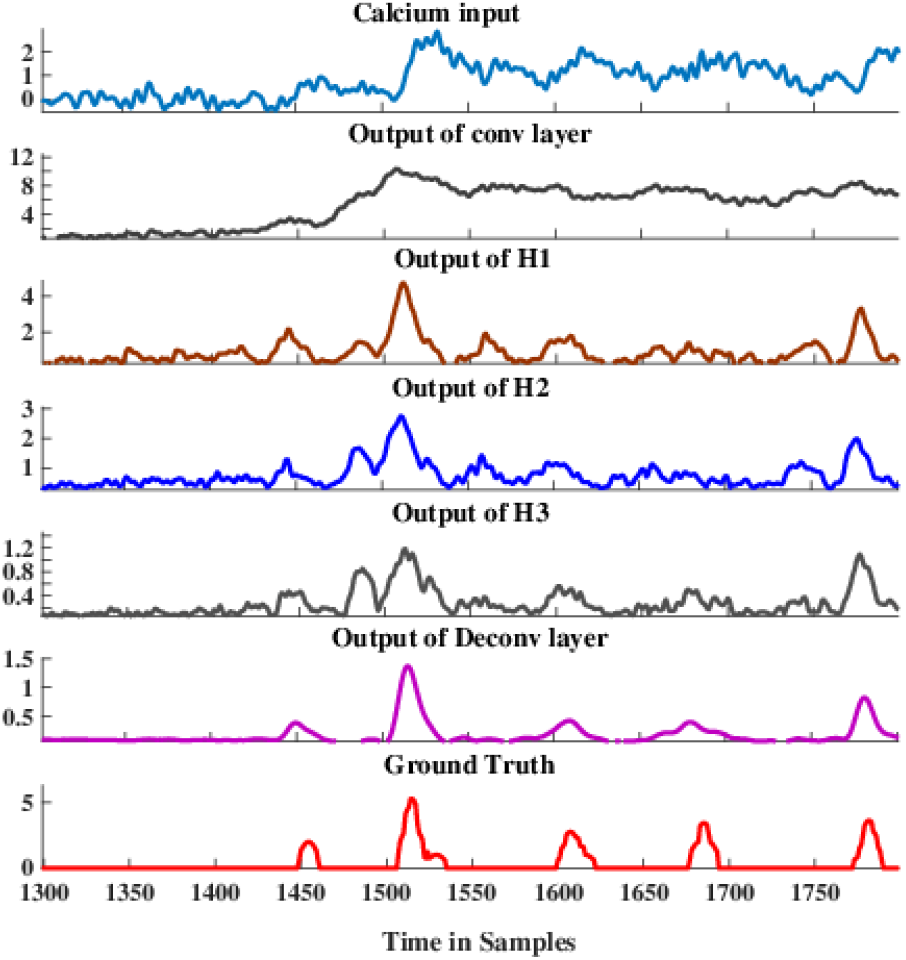
Layer-wise output of a 3-dense layer S2S. Please note that small calcium changes which are not clearly distinguishable in the original fluorescence signal are amplified in the output of the deconvolution layer.

#### 2.4.5 Filter Responses

The S2S method learns to synthesize the spike information from the calcium fluorescence signals due to the filters in the analysis and synthesis convolutional layers. Hence, we analyzed the frequency response of the filters. Fig 7 shows the cumulative frequency response of the filters in the analysis convolutional layer and the synthesis convolutional layer. It can be observed that the analysis layer gives emphasis to low-frequency information (between 0-5 Hz). In the case of the synthesis layer, we observe a harmonic structure, which could be attributed to the fact that the layer predicts the spike signal. The harmonic structure is present in the time-domain analysis as well. This is probably because the network is learned to predict spike trains with different time-scales and frequency components. The de-convolution layer that synthesizes the signal at the output makes our approach distinct. It is more effective than the neural network-based methods that estimate spikes in a sample-by-sample manner (please refer to Section 2.3).

**Fig 7.**
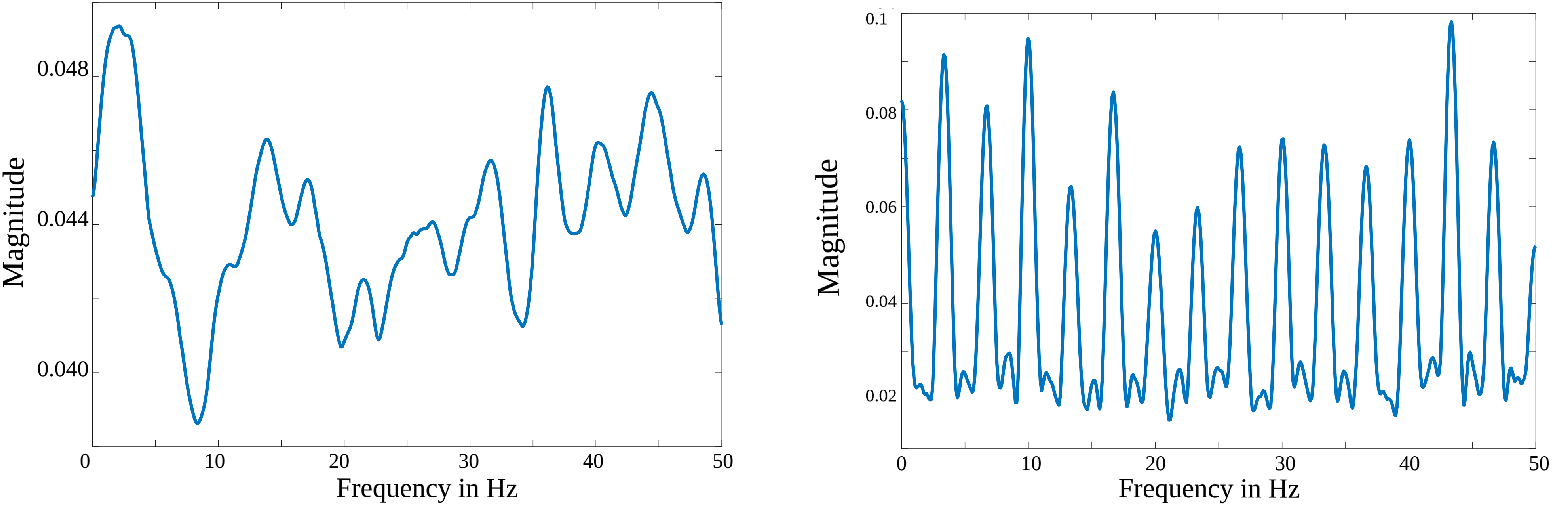
Cumulative frequency responses of filters: (***left***) Shows the frequency response of the analysis filters. (***right***) Shows the response of synthesis filters having larger amplitudes at equal frequency intervals. The energy at the output of S2S is purely concentrated on spikes both at the temporal and the spectral domain.

## 3 Discussion

We showed that the proposed S2S method can yield state-of-the-art results in the spike estimation task. Our method resulted in significant improvement in both primary and secondary evaluation measures. This architecture seems to be appropriate for the spike estimation problem. The synthesis layer (which reconstructs the spike signal) and the cost function based on the correlation coefficient resulted in faster convergence and better performance.

### 3.1 Nature of Improvement

We discuss the nature of the improvement achieved by S2S method. S2S had significantly better linear and non-linear correlation measures and a comparable AUC value to that of the baseline algorithms. This is reflected in the spike estimate by having an overall shape and the resolutions of the individual estimates similar to the ground truth spike train at the 40 ms resolution. We further observe that these differences are preserved at a higher resolution (10 ms bin width) as well (Please refer to the scalability experiments in Section 2.3). Hence, S2S is able to infer spike trains (100 Hz) as well as the firing rates (25 Hz). Fig 8 compares the spike estimates obtained from the S2S and the baseline convi6 method to the discrete ground truth at 40 ms bin width.

**Fig 8.**
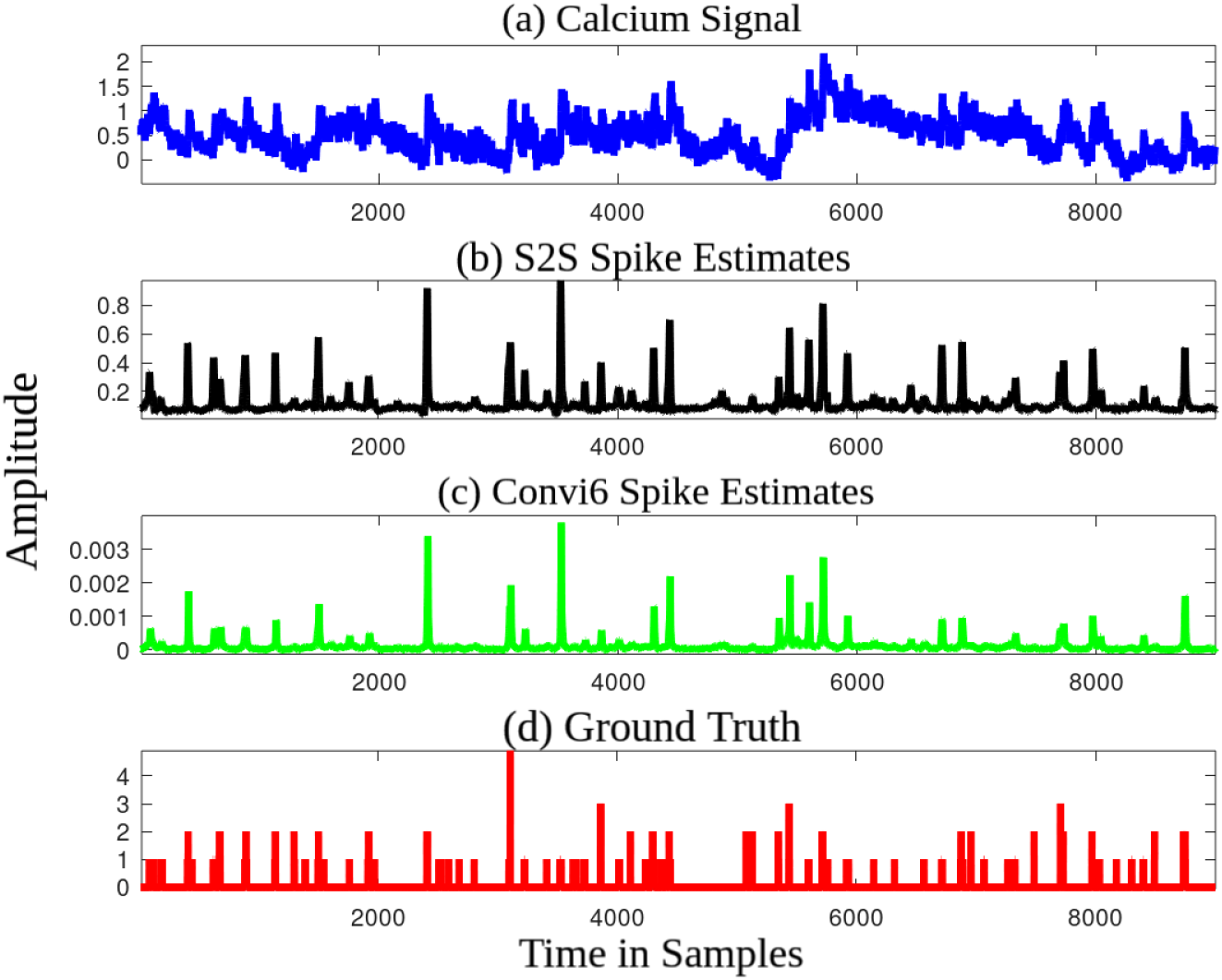
Comparison of spike estimates. (a) An example calcium input signal, spike estimates of (b) S2S and (c) convi6 methods, and (d) discrete ground truth. Observe the similarity in shape between the S2S and the ground truth, compared to the convi6 method.

It can be observed that the peak locations are more resolved in S2S than the convi6. The dynamic range of S2S is much larger than the baseline, and this is reflected by the correlation measure. For every action potential, the convi6 has a very small increase in the amplitude of the estimated signal, typically around 0.001. The corresponding increase in the amplitude of S2S spike estimate is 0.2-0.3. The amplitude of the individual S2S spike estimates are especially high when the actual spike count is greater than one or when a burst of spike occurs, similar to the ground truth. The amplitude of the convi6 estimates are shorter than that of the S2S estimates.

To confirm that, we computed the average deviation of the amplitude between the spike estimate and the ground truth. The average deviation of the S2S spike estimate with respect to the ground truth is much smaller than that of the convi6 method. The average deviation varies from 0.02 to 0.45 per test file for the S2S spike estimate, whereas it varies from 0.55 to 1.48 per test file for the convi6 spike estimate. To further validate the statistical significance of the deviation per test file, we conducted a pair-wise t-test considering all the 32 test files. The null hypothesis (H) was that there is no statistical significance in the deviation per test file. The null hypothesis is rejected with a very high confidence(*α* = 3.1E-30). We also observed that the improvement in performance of S2S method is independent of Gaussian windowing that is applied optionally to the discrete spike estimates during the training.

In Fig 8, it can be observed that the relative spike positions at the output of S2S are similar to that of convi6, as also pointed out by the AUC measure. Hence, the improvement in linear and non-linear correlation measures can be mainly attributed to the improved detection of small calcium events which result in a spike.

### 3.2 Design Aspects of the S2S Method

We examined various design aspects of the S2S method. Our studies revealed that: (a) making the targets less sparse by convolving the ground truth discrete spike signal with Gaussian window helps, (b) at least one hidden layer is needed to resolve the spike locations, (c) the method is not sensitive to initialization; converges within 50 epochs, and yields similar performances when retrained (the results are reproducible), and (d) the method is capable of generalizing across unseen indicators.

One of the major concerns in using a supervised approach for spike inference is its computational complexity. Generative approaches, on the other hand, require mainly novel parametric settings for new datasets. S2S method, although being a supervised method, is computationally efficient. All the top-performing supervised algorithms in the spikefinder contest had recurrent units in the architecture, which resulted in increased training time. The proposed S2S sytem is 50 times faster than the best performing supervised model when trained with GPU using Sun Grid Engine, potentially because the network has very few trainable parameters and the use of time-distributed hidden layers. As presented in the Methods section, the three hidden layer S2S has only 8790 trainable parameters.

### 3.3 Multiple Evaluation Measures

Performance evaluation of a spike estimation algorithm should consider the spike estimate’s overall shape, monotonic relationship with the ground truth, and the estimates’ accuracy with multiple thresholds. There is no unique measure that encompasses all this information [34]. For instance, the spike train is considered as a density function, and its shape is evaluated with the Pearson’s correlation coefficient. However, this measure is invariant under affine transformations. Thus, it is not easy to interpret the outputs as spike counts or rates. The Spearman’s rank-order correlation coefficient evaluates both the strength and the direction of the association between the two ranked variables. The dynamic ranges of the spike information and discrete spike train are employed for ranking. Unlike the Pearson’s correlation coefficient, this measure considers the non-linear relationship between the variables. Finally, the area under the receiver operating characteristics (AUC) measures how well the spikes have been detected. AUC is not sensitive to changes in the relative height of different parts of the spike information. Thus, this measure alone is not adequate. The proposed S2S method outperforms all the other systems in terms of Pearson’s correlation coefficient and Spearman’s rank correlation coefficient and achieves similar performance in AUC (only the DeepSpike method has marginally high AUC). This indicates that the S2S is yielding a better estimate of spike signal than existing generative and supervised methods. Specifically, the S2S method resulted in an improvement of 46% in primary evaluation measure compared to the STM baseline, which is two times that achieved by the Spikefinder contest (23%). This performance gain could help in further directions in spike estimation based on signal reconstruction strategies.

## 4 Methods

### 4.1 Signal-to-signal Neural Network for Spike Signal Estimation

As discussed earlier in the Introduction section (Section 1), approaches are emerging to convert one sequence into another sequence in an end-to-end manner in various fields related to sequence processing such as speech processing and natural language processing. For instance, converting a sequence of acoustic features to a sequence of letters or words and translating the sequence of words from one language to another language. We cast the spike estimation problem from the calcium fluorescence signal as a sequence-to-sequence conversion problem, as a signal is a time-ordered measurement. Towards that, we take inspiration from recent work on end-to-end speech source separation [25] to a develop signal-to-signal network which learns to predict/estimate spike signal given the calcium fluorescence signal as input (as illustrated in Figure S1). Intuitively, the S2S method can be seen as a single-channel signal enhancement system where the desired spike signal present in the calcium fluorescence signal is enhanced, while the undesirable signals or variabilities are suppressed.

#### 4.1.1 S2S Architecture

The S2S method consists of a convolution layer followed by a fully connected dense layer(s) and an output convolution layer. The hidden layers are implemented in a time-distributed manner, i.e., the output of the first convolution layer for every segment of the calcium fluorescence signal is independently processed by the hidden layer(s) and then synthesized by the output convolution layer. In the network, the output of each hidden node or filter is fed to a rectified linear unit (ReLU) non-linearity, except for the output convolution layer. ReLU units in the neural network do not saturate unlike *sigmoid* and *hyperbolic tangent* activations. They also help in learning a non-negative representation in the successive layers.

The hyper-parameters of the S2S system are: the length of signal input *w*_*seq*_, kernel width *kW*_*in*_ and kernel shift *dW*_*in*_ of the input convolution layer, number of filters in the input convolution layer *nFilt*_*in*_, number of hidden layers *I* and the number of nodes *nhu*_*i*_ in each hidden layer *i* ∈ {1, . . . *I*}, kernel width *kW*_*out*_ and kernel shift *dW*_*out*_ of the output convolution layer and the number of filters in the output convolution layer *nFilt*_*out*_. Based on the systems reported in the spikefinder challenge, we set *w*_*seq*_ = 100 samples (i.e., 1 sec). We set *kW*_*in*_ = *w*_*seq*_ and *dW*_*in*_ = 1 sample. In the output convolution layer, *kW*_*out*_ = *nhu*_*I*_ (i.e. number of nodes in the last hidden layer), *dW*_*out*_ = 1 sample and *nFilt*_*out*_ = *w*_*seq*_. In other words, for every frame of 100 sample input, the output convolution layer synthesizes 100 samples of spike signal, which are then overlapped and added to get a spike signal of same length as that of the input calcium fluorescence recording. The number of frames is determined by *dW*_*in*_, which is one sample. The remainder of the hyper-parameters *nFilt*_*in*_ = 30, *I* = 3 and *nhu*_1_ = *nhu*_2_ = *nhu*_3_ = 30 were determined by cross validation on the training set through a coarse grid search. Thus, the three hidden layer S2S compared against the state-of-the-art methods in the Results section has (100 × 30 + 30) + (30 × 30 + 30) + (30 × 30 + 30) + (30 × 30 + 30) + (30 × 100) = 8790 parameters. In the case of the one hidden layer study presented in the Analysis section, the network has (100 × 30 + 30) + (30 × 30 + 30) + (30 × 100) = 6960 parameters, and in the case of no hidden layer (100 × 30 + 30) + (30 × 30) = 6030 parameters.

#### 4.1.2 Training the S2S System

We train the S2S system by maximizing the Pearson’s correlation coefficient between the spike signal estimated by the network and the ground truth spike signal. We also investigated other cost functions such as mean square error and cross-correlation with sigmoid non-linearity. The Pearson’s correlation coefficient yielded the best system on both the cross-validation experiments conducted on the training set and the test set. For training the S2S, we split the training data in the spikefinder challenge into two parts: a training set (80%) and a validation set (20%). At each training epoch, the training set was used for training the network parameters, and the validation set was used for cross-validating the network. We used Adam optimizer with a starting learning rate of 0.001 and a batch size of 20. Early stopping with a patience factor of 6 is used to stop the training whenever the validation loss increases compared to the previous epoch. As the ground truth is a sparse discrete signal, the target spike signal was optionally convolved with a Gaussian window to make the targets less sparse for the network’s efficient training. The Gaussian window (11, 5) was obtained through cross-validation, i.e., the window that yielded the best validation cost. The implementation was carried out on Keras [43] with TensorFlow [44] backend. Upon acceptance of this paper, the software will be made available at https://github.com/Jiltseb/S2S_for_spike_inference.

### 4.2 Data

We used the spikefinder challenge dataset to validate the proposed S2S method. We followed spikefinder evaluation [15] owing to the following reasons: First, it is one of the largest publicly available dataset containing different scanning rates and methods and calcium indicators. Second, the spikefinder challenge provides a bench-marking framework to compare different spike estimation methods using different evaluation measures. Finally, the challenge provides state-of-the-art baseline systems to which the proposed method can be compared to. As mentioned earlier, the spikefinder dataset has five bench-marking datasets consisting of 92 recordings from 73 neurons. Five additional datasets from GENIE project [41] have been provided to ensure that the supervised models do not over-fit to the training data and to test the generalization ability. The zoom factor of all the recordings is at use-case resolutions for the calcium imaging experiments. For further information, the reader is referred to [15]. The dataset is available at the open-source, public repository (https://github.com/codeneuro/spikefinder).

### 4.3 Comparison Framework

The proposed method’s performance was compared to two sets of baselines: The results were first compared to the top-five algorithms in the spike finder contest. They are either generative [11, 32] or supervised [15, 17, 33, 39] approaches. The generative approach, which was initially published in [11] uses a biophysical model and estimates the maximum probable spike information from the fluorescence signals. Efficient Bayesian inference is performed on this MLSpike model, including a drifting baseline and nonlinear modeling of calcium to fluorescence conversion. MLspike was the winner of the spikefinder contest. This algorithm’s performance is boosted (w.r.t. the original work) owing to the parameter tuning with respect to the challenge dataset. The second best generative approach [32] was based on an auto-regressive approximation to the calcium fluorescence signal. Spike information was then estimated by solving a non-negative sparse optimization problem. Based on the advances in neural network-based models for various applications, it is not surprising that 7 out of top-10 algorithms are deep learning-based supervised algorithms (see Table 1). All the top-performing algorithms used recurrent layer in the network. For the best-performing algorithms, we have taken the evaluation results from the spikefinder challenge [15], and by running the evaluation scripts (https://github.com/berenslab/spikefinder_analysis).

We compared the proposed network’s performance, and efficiency with the best performing supervised baseline, available as an open-source Python software (https://github.com/kleskjr/spikefinder-solution). We trained the network using open-source software. However, the model weights provided by the “convi6” authors yielded better linear correlation values than those obtained by training the neural network. Convolutional layers take a broad temporal context to predict a single output corresponding to the spike information. The dataset index that was provided as an auxiliary input to the network observed to improve the system’s performance. Convolutional layers are followed by an intermediate LSTM-recurrent layer and further by convolutional layers of smaller width. The learned filters collectively capture the spike-related information.

### 4.4 Evaluation

All the evaluations were done at 25 Hz (40 ms bin width), following the protocol used in spikefinder evaluations (https://github.com/berenslab/spikefinder_analysis). As per the protocol, we used the Pearson’s correlation coefficient as the primary evaluation measure. Δ correlation is computed as the average difference in correlation coefficient compared to the STM algorithm (Please refer to Table 2). The secondary evaluation measure was Spearman’s rank correlation coefficient, which considers the direction and non-linear relationship between the estimates and actual spikes. The area under the ROC curve (AUC) curve was the final measure, which evaluates spikes’ detection in a 40 ms bin. We used the function *roc_curve* from *scikit-learn* [45] Python package for computing the AUC and *spearmanr* function from *scipy* [46] Python package for computing the rank. These evaluation measures collectively denote the similarity with the ground truth. We consider them in the order of their preference as in the spikefinder challenge. The test set’s rank measure was not included in the challenge paper, although it was the secondary measure on the training sets.

## 5 Conclusion

This paper presented a neural network-based signal-to-signal conversion approach (S2S) for spike estimation from imaging data. The neural network analyzes and filters the raw calcium imaging data at the input and synthesizes or estimates the spike signal at the output, in an end-to-end manner. During training, the parameters of the neural network are estimated by a cost function based on Pearson’s correlation coefficient between the estimated spike signal and the ground the truth. An evaluation on the Spikefinder benchmarking data set showed that the proposed approach achieves state-of-the-art performance in the spike estimation task. We discussed statistical significance of the results, scalability of the results at a higher sampling rate, generalization ability of the S2S method and analyzed the layer-wise outputs and the convolutional filter responses. We further commented on the effect of Gaussian training target and the number of hidden layers in the architecture.

## Acknowledgements

This work was partially supported by the Swiss Government Excellence Scholarship Project with ESKAS No: 2017.0575. The authors would like to acknowledge the contribution of Mr. Pavan Kumar Dubagunta for his fruitful discussions towards the work and Mr.Mari Ganesh Kumar for his review of the draft.

## Footnotes

1 In this article, we use the terms dense layer and hidden layer interchangeably.

## Notes

### Competing Interest Statement

The authors have declared no competing interest.

